# Embryo-scale epithelial buckling forms a propagating furrow that initiates gastrulation

**DOI:** 10.1101/2021.12.14.472566

**Authors:** Julien Fierling, Alphy John, Barthélémy Delorme, Alexandre Torzynski, Guy Blanchard, Claire Lye, Grégoire Malandain, Bénédicte Sanson, Jocelyn Étienne, Philippe Marmottant, Catherine Quilliet, Matteo Rauzi

**Author notes:** Contributed equally (Julien Fierling, Alphy John, Barthélémy Delorme, Jocelyn Étienne, Philippe Marmottant, Catherine Quilliet, Matteo Rauzi). To whom correspondence should be sent, and.

## Abstract

Cell apical constriction driven by actomyosin contraction forces is a conserved mechanism during tissue folding in embryo development. While much effort has been made to better understand the molecular mechanisms responsible for apical constriction, it is still not clear if apical actomyosin contraction forces are necessary or sufficient *per se* to drive tissue folding. To tackle this question, we use the *Drosophila* embryo model system that forms a furrow on the ventral side, initiating mesoderm internalization. Past computational models support the idea that cell apical contraction forces may not be sufficient and that active or passive cell apico-basal forces may be necessary to drive cell wedging and tissue furrowing. By using 3D computational modelling and *in toto* embryo image analysis and manipulation, we now challenge this idea and show that embryo-scale force balance of the tissue surface, rather than cell-autonomous shape changes, is necessary and sufficient to drive a buckling of the epithelial surface forming a furrow which propagates and initiates embryo gastrulation.

## 1 Introduction

The spectacular epithelial coordination that characterizes morphogenesis in the course of embryo development involves long-range cell interaction. Previous studies support the idea that long-range tissue mechanics direct morphogenetic coordination [1, 2, 3, 4, 5, 6, 7]. Here we investigate the role of embryo-scale mechanics in forming a furrow in an epithelial tissue. Furrow formation is a process that involves the bending of a tissue along a line. The formation of a furrow is pivotal during embryo development since it initiates tissue topology changes structuring the future animal. Unravelling the mechanisms which drive furrowing is thus key to understanding how vital processes such as gastrulation and neurulation are initiated. Furrow formation, which results in tissue folding and eventual internalization, is a tissue shape change that emerges from a collective cell behaviour. We thus need to understand how cells mechanically interact and collectively drive tissue furrowing. Ventral furrow formation (VFF) during early gastrulation in the *Drosophila* embryo is a well studied process: a fold along a line at the ventral side of the embryo (the prospective mesoderm) forms parallel to the anterior-posterior direction [8, 9]. Numerous *in vivo* studies have tackled the mechanisms driving VFF. Cell apical constriction [10], basal expansion [11], lateral tension [12,13] and cytoplasmic flow [14] are mechanisms that have been studied and that are associated with cell shape changes (e.g., from columnar to wedge-shaped) and the eventual formation of the furrow. Cell apical constriction driven by the contraction of apical actomyosin networks is one of the most thoroughly investigated mechanisms since it is a striking change in shape, given its magnitude and rapidity, and it is amenable to analysis since it occurs at the surface of the tissue where microscope-based imaging technologies can be more easily applied. While numerous studies have highlighted the molecular nature and the key role of apical constriction in driving furrow formation [10,15,16,17,18], it is still not clear if this is necessary or sufficient *perse* to form the ventral furrow in the gastrulating *Drosophila* embryo. Computational modelling is a powerful tool to study the mechanics and the change in shape of tissues: minimal models recapitulating key features of an epithelium can be engineered and limitlessly tested to investigate the fundamental mechanisms driving a process. 3D and 2D cross-section models based on active deformation and active stress, respectively, have been developed to understand the mechanics of VFF [19]. Due to simplifying hypotheses, the former cannot inform us on the stresses at work [20, 21], whereas the latter lack the 3D geometry necessary to incorporate the global force balance within which the furrowing process takes place [22, 23, 24, 25, 13, 26]. 2D models, mimicking the embryo crosssection, have been used to predict that apical contractions may not be sufficient to drive furrow formation. By implementing an experimental strategy combining computational modelling, infrared femtosecond (IR fs) laser manipulation coupled to multi-view light sheet microscopy and quantitative image analysis, we show that apical contraction forces are necessary to drive VFF. In addition, we unveil a new emergent long-range mechanism based solely on surface mechanics (i.e., free of cell lateral and basal forces and not relying on cell shape changes from columnar to wedged) over a curved tissue that is sufficient to drive furrow formation.

With the aim of setting common grounds, we provide in Box 1 the definition of key terms that will be extensively used in this manuscript.

#### Box 1. Definition of key terms

**Furrowing:** the process by which a tissue bends forming a fold having much greater bending curvature along one direction compared to the orthogonal direction. **Invagination:** the process by which a tissue is internalized inside the embryo. **Buckling:** a sudden change in the type of mechanical equilibrium, from a state in which load is mostly balanced by internal forces along the plane of the tissue to a state in which load is mostly balanced by internal forces normal to the tissue. **Strain:** the magnitude of deformation of material having a given shape (called the current configuration) at a specific time and location with respect to its equilibrium configuration. In the absence of pre-strain, the equilibrium configuration is the original shape. **Pre-strain:** strain associated with a change of the equilibrium configuration rather than changes of the current configuration. For instance, an increase of the activity of the molecular motor MyoII corresponds to an increase in pre-strain, not necessarily entailing a change of shape (current configuration) with respect to the original shape. **Stress:** the internal tension within the material which is elicited by strain. **Pre-stress:** stress which is due to a change of the equilibrium configuration (i.e., the pre-strain).

**Boxed figure 1.**
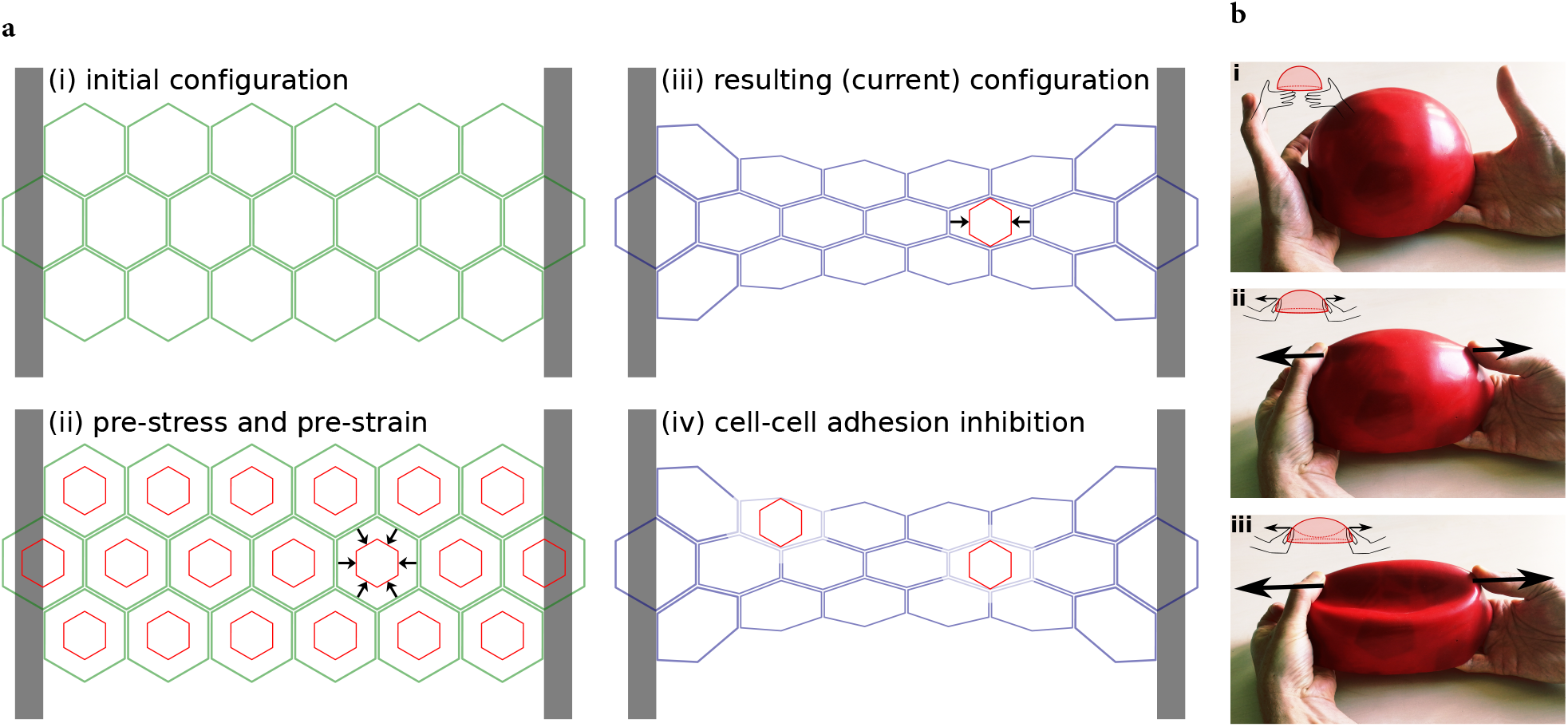
(**a**) Deformations and stresses in a tissue with anisotropic boundary conditions and isotropic actomyosin contractile activity. (i) Initial configuration of a tissue held between two clamped boundaries at its longitudinal ends and free at its latitudinal ends. Without MyoII activity, this represents the equilibrium configuration. (ii) Representation of the effect of uniform isotropic MyoII activity. The initial configuration (green) is no longer the equilibrium configuration: the red shape shows the equilibrium configuration of each cell, which is thus pre-strained in its initial configuration. Black arrows indicate the corresponding pre-stress (here shown for one cell). (iii) Current configuration (blue) resulting from MyoII pre-stress. Tissue cohesion and fixed boundary conditions prevent cells from adopting their equilibrium configuration (shown in red for one cell). Along the latitudinal direction, away from the boundaries, cells are able to contract to their equilibrium size, thus strain (nearly) equals pre-strain in this direction, which results in (close to) zero stress along this direction. Along the longitudinal direction, tissue cohesion and boundary conditions prevent any length change compared to the initial shape, thus strain is zero and stress (arrows) is equal to pre-stress along this direction. (iv) If cell-cell cohesion is inhibited, cells relax to their equilibrium shape (red), revealing the anisotropy of stress in the configuration (iii) [27, 28]. (**b**) An example of buckling of an elastic sheet. (i) A thin elastic shell in its equilibrium configuration. (ii) Forces applied in two positions deform the elastic shell. Note the flattening (reduction of curvature) and the elongation (in-plane deformation). (iii) Above a threshold level of applied force, buckling occurs. Note the depression (inversion of the sign of curvature) surrounded by a highly curved rim. The change of shape happens suddenly.

## 2 Results

### 2.1 Apical contraction is necessary to drive furrow formation

A large body of genetic perturbation experiments, resulting in eventual cell apical constriction and mesoderm internalization failure, support the idea that apical contraction of actomyosin networks is necessary to drive tissue furrowing [27, 30, 31, 32, 33, 34]. In order to test the role of actomyosin contraction forces in mesoderm invagination, Guiglielmi and colleagues in 2015 developed a two-photon optogenetic technique that allows to indirectly perturb the actin cytoskeleton by modulating with spatial and temporal specificity the concentration of plasma membrane phosphoinositides (PIP2) that also plays a role in G-actin polymerization [35, 36, 37, 38, 39]. This study shows that depletion of cortical PIP2 at the cell cortex of mesoderm cells inhibits apical constriction and mesoderm invagination. We now aimed to test more directly and specifically the role of apical contraction forces in mesoderm furrow formation. To that end, we implement IR fs laser dissection, a technique shown to sever the actomyosin network with high spatio-temporal specificity without compromising cell membrane integrity [40, 41]. After laser dissection of the ventral actomyosin cytoskeleton, the network recoils and the cell apical surface dilates. The actomyosin network eventually recovers restoring apical contraction forces and cell apical constriction (Fig. 1a and Supplementary movie 1). IR fs ablation is therefore an effective tool to sever contractile actomyosin networks while preserving cell integrity. We performed ablation of the contractile actomyosin network during furrow formation (i.e., when the ventral tissue has a concave shape) and monitored tissue curvature changes. After ablation, the ventral tissue changes curvature from concave to convex. Eventually, after actomyosin network recovery, the ventral furrow regains its concave shape (Fig. 1c and b). This directly demonstrates that apical contraction forces, driven by apical actomyosin networks, are necessary for furrow formation. To test whether apical contraction forces are also necessary for tissue invagination, we performed sequential laser dissections of the ventral actomyosin network to repeatedly downregulate actomyosin contraction forces that would otherwise recover within 10 seconds (Fig. 1a and Supplementary movie 1). Under these conditions, ventral cells remain at the embryo surface resulting in tissue invagination failure (Supplementary movie 2). This demonstrates that apical contraction forces generated by actomyosin networks are necessary for both furrow formation and subsequent tissue invagination.

**Figure 1:**
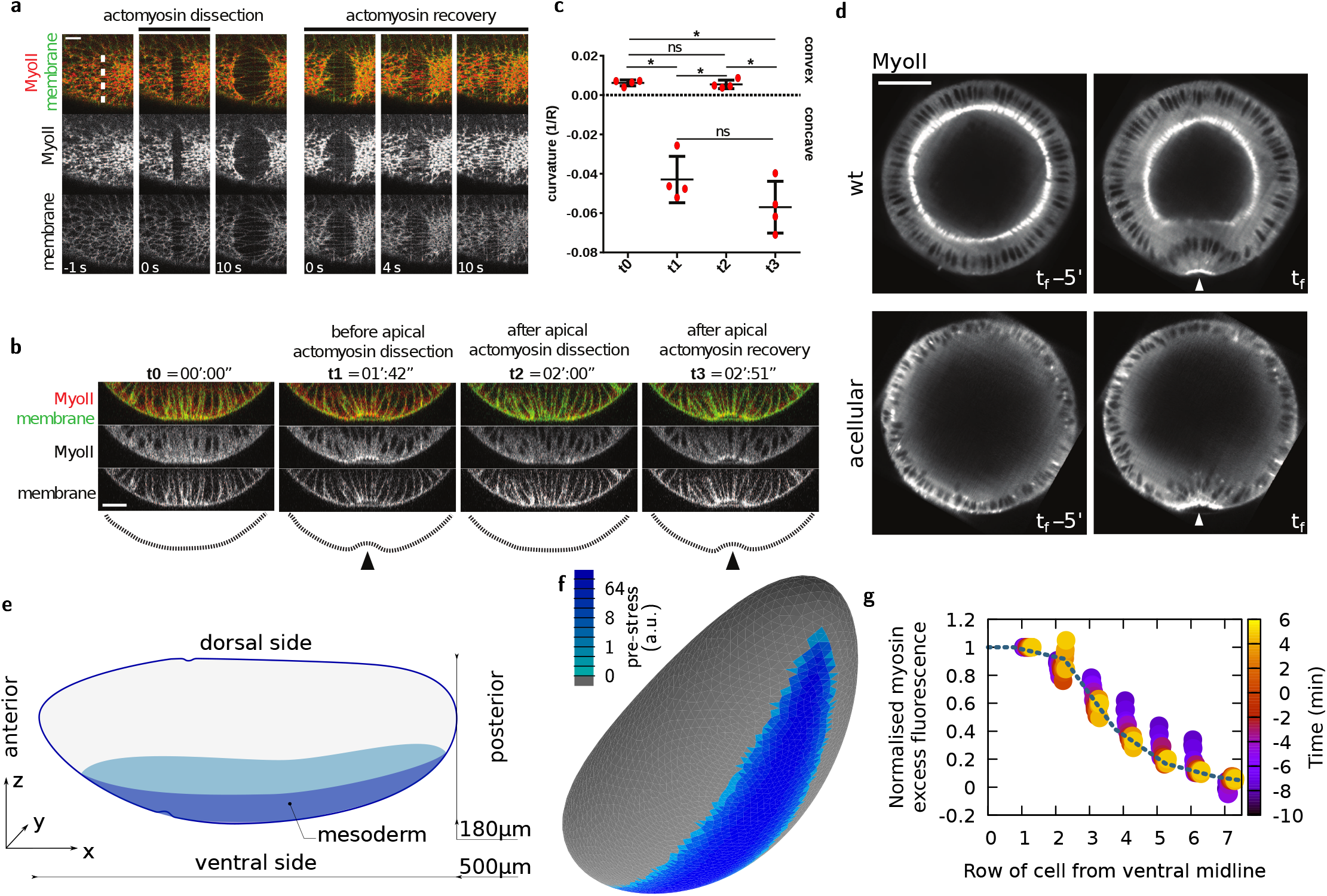
Using IR fs laser dissection to demonstrate the necessity and building of a model to test the sufficiency of apical contraction to drive VFF. (**a**) Recoil and recovery of the ventral apical actomyosin network after laser dissection. Scale bar 10 μm. (**b**) Cross-sectional view of the embryo ventral side just before (t0 and t1) and just after (t2) laser dissection, and (t3) after actomyosin recovery. Scale bar 20 μm. (**c**) Curvature analysis before and after laser ablation and during actomyosin network recovery as shown in **b.** (**d**) Digital mid-cross-sections before and during furrow formation in wild-type and *slam*^-^ *dunk*^-^ acellular embryos. Scale bar 50 μm. (**e**) Representation of the embryo geometry. (**f**) Finite element mesh of an embryo-shaped elastic surface where colour-marked facets are pre-strained to mimic MyoII activity (log scale). (**g**) Circles, normalized MyoII profile at different phases of VFF as a function of distance from the ventral midline in experiments (average of *n* = 3 embryos). Line, pre-stress profile chosen for simulations. This profile is also similar to the one reported by [29].

### 2.2 Inferring sufficient conditions to form a furrow in an active ellipsoidal model surface

During VFF, the *Drosophila* embryo is constituted of a single layer epithelium with the cell apices facing outwards (see Fig. 1d). In embryos carrying mutations for the genes *slam* and *dunk (slam^-^dunk^-^),* the process of cellularization (prior to VFF) is impaired: the apical membrane fails to ingress to form lateral and basal side of blastoderm cells, resulting in acellular embryos [14]. In acellular embryos, nuclei of mesoderm cells still displace basally [14] and a ventral furrow still forms (Fig. 1d). Therefore, we asked whether apical forces alone lead to deformations similar to those observed *in vivo.* We modelled the apical surface of the *Drosophila* blastoderm as a thin elastic surface (Fig. 1e). A more general approach would have been to consider a visco-elastic model, since the relaxation time of the *Drosophila* embryonic epithelium is estimated to be around one minute [42], shorter than the process of VFF [4]. However, a visco-elastic model would have involved both a much higher computational complexity and a much greater difficulty of exploring the parameter space to fit the data. Since the mechanical load due to molecular motor Myosin II (MyoII) activity is constantly increasing during the early phase of the VFF process [29], we hypothesised that the effects of the viscous relaxation would be negligible in comparison to the elastic response to the increased load. Under this hypothesis, a purely elastic model is sufficient to recapitulate the main features of this fast process.

We modelled the initial three-dimensional shape of the *Drosophila* embryo as in [5] (Supplementary Information). The global shape of the embryo is ovoid, with the long axis (along anteroposterior, AP) three times the length of the short axis (along dorsoventral, DV). In good approximation to live embryos, our model presents a dissymmetry with respect to the mid-coronal plane, the ventral side being more curved than the dorsal, while it is symmetric with respect to the mid-transverse and mid-sagittal planes. As in previous models, the volume within this elastic surface is assumed to be constant throughout the simulations. The embryo shape is also constrained by the vitelline membrane, which is undeformable, and separated from it by a thin layer of perivitelline fluid. We assumed that, before VFF, the distance between cell apices and the vitelline membrane is approximately 0.2 - 0.5 μm. This distance is allowed to vary locally [43, 23, 44, 45] while the global volume of perivitelline fluid is kept constant. We assume that tangential forces, such as friction or specific adhesion between the apical elastic surface and the vitelline membrane, are negligible at this stage, consistent with experimental and theoretical results [44, 45, 5]. We also hypothesise that viscous shear forces exerted on the surface are negligible.

Using the finite element software *Surface Evolver* [46], we model the mechanics of the apical surface of the embryo as a thin elastic shell of given elastic properties [47]. In this continuous model, the tissue is not partitioned into cells, nevertheless regions of space can still be assigned different properties, reflecting distinct portions of the embryo showing different gene expression patterns and mechanical properties [19, 5]. Therefore, we assign a local active pre-stress to finite element facets in the region corresponding to the embryo prospective mesoderm (see Fig. 1f). We do so by assigning these facets a target area *A*_0_ which is smaller than their initial area *A_i_*, Fig. 2a. This means that, before deformations cause relaxation, these facets experience pre-strain 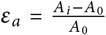 and thus are pre-stressed by 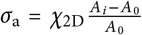, where *χ*_2*D*_ is the in-plane compression elastic modulus (see Supplementary Information and Supplementary Fig. 1a). We find that the pre-stress applied in the mesoderm region results in increased tension not only at the ventral but also at the lateral and dorsal regions of the embryo, Fig. 2b. For an explanatory definition of pre-strain and pre-stress, see Box 1.

**Figure 2:**
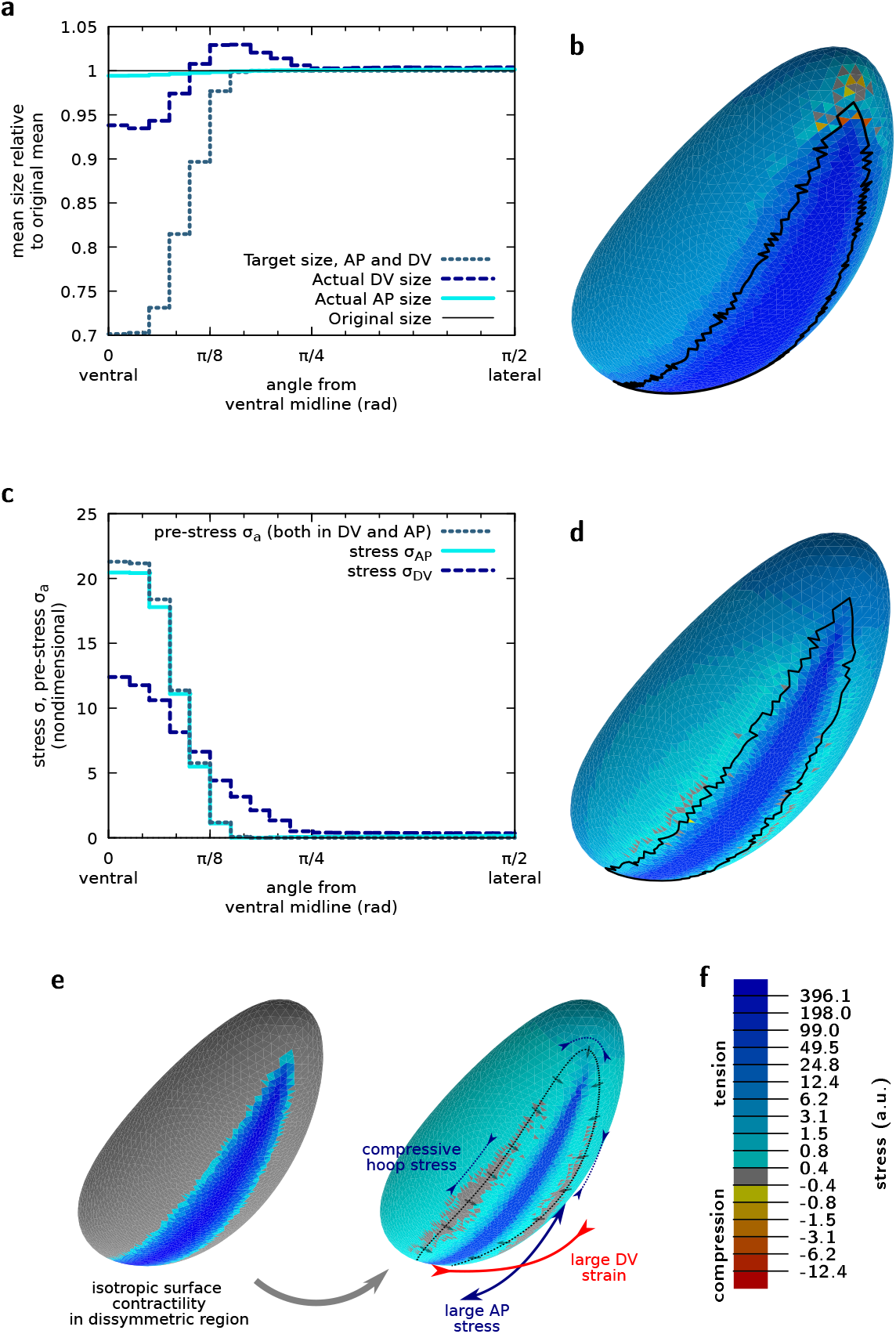
Actomyosin contractility drives tension anisotropy on the ventral side of the embryo. (**a**) Strain angular profile (current size relative to initial size) along the AP and DV axes for midline pre-strain 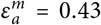. (**b**) Mechanical stress (sum of the two principal stresses) resulting from the area pre-strain. Midline pre-strain 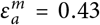. Black line corresponds to the boundary of the pre-strained region in the current configuration. See tensor components in Supplementary Fig. 1b. (**c**) Angular profiles of the pre-stress *σ_a_* and of the two principal stresses along the AP and DV axes for 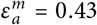. (**e**) Isotropic pre-strain pattern (left) yields anisotropic mechanical response, with a greater stress and strain along the AP and DV axes, respectively. The cells at the periphery of the mesoderm move towards it, arrows, which generates a hoop stress along the dotted line. (**d**) Same as **b** for 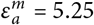. See tensor components in Supplementary Fig. 1d and profiles in Supplementary Fig. 1e and `f`. (**f**) Colour code for panels **b** and **d.** All panels are for nondimensional mechanical parameters 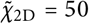 and *ν*_2*D*_ = 0, see Supplementary Information.

It has been observed that MyoII activity is not uniform across the mesodermal cells: MyoII shows a graded distribution over approximately seven AP rows of cells on both sides of the mesoderm with highest level at the mid-line [29]. Experimentally, we have found that this spatial profile is well preserved during VFF (see Fig. 1g). We have thus implemented a graded distribution of pre-stress in our computational model mimicking the graded MyoII distribution. To this end, we have applied a gradient of pre-strain which starts from the ventral mid-line where *ε_a_* is equal to 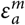 and decreases according to the profile shown in Fig. 1g. The resulting stress is largest in the region where the pre-stress is applied, but decreases laterally with a profile that differs from the one of the pre-stress gradient, Fig. 2b.

### 2.3 Tension anisotropy emerges from tissue and embryo geometry

The increase in tension in the mesoderm during *Drosophila* VFF is known to be anisotropic [27], with a higher tension along the AP axis than along the DV axis. It has been shown that tension anisotropy is the cause of the anisotropic organisation of the actomyosin network that eventually arises during VFF [28] and it has been suggested that the geometry of the embryo is responsible for the emergence of this anisotropy of tension [48, 28]. We thus considered whether a mechanical model would predict anisotropic tension by imposing isotropic MyoII activity. It has been shown that in an elastic flat plate, isotropic active pre-stress in an asymmetric geometry causes stress and strain anisotropy, which are respectively larger along the long and short axis of the domain [49,50]. Our simulations show that this effect is also at play in the embryo 3D geometry, and that mechanical tension is strongly anisotropic in the mesoderm, AP tension being twice the DV tension along the ventral midline (Fig. 2c and e and Supplementary Fig. 1a and b).

What is the origin of the stress anisotropy along the AP and DV axes? By decomposing the length of the contracting ventral tissue along AP and DV, it is noticeable that this length is about three and six times less than the total blastoderm length along the mid-sagittal and mid-cross sections, respectively (Supplementary Fig. 1c). Therefore, the stretch of neighbouring tissue which is necessary to achieve the same contraction strain along the two orthogonal directions is about three times greater along the AP than the DV axis. This results in a three times greater cell resistance to stretch along AP than DV. The shape anisotropy of the system would thus explain why the surrounding tissue appears more difficult to deform along the AP than along the DV direction even though mechanical properties of the entire tissue surrounding the mesoderm are imposed to be the same.

More specifically, the AP stress σ_AP_ is found approximately equal to the pre-stress σ_a_ initially imposed on the ventral region. This is consistent with the fact that little tissue deformation takes place along the AP axis (Fig. 2c), and can be related to the mechanical behaviour of contracting actomyosin networks anchored to stiff boundaries which oppose resistance to deformation [51]. On the other hand, the DV stress σ_DV_ is found to be different from the pre-stress, both in the ventral and in the ventrolateral region. This is consistent with the differential tissue deformation taking place along the DV direction: in the highly contractile ventral region (for angles between 0 and *π*/8) the tissue contracts (Fig. 2a) resulting in σ_DV_ smaller than σ_a_ (Fig. 2c). In the ventro-lateral region (for angles between *π*/8 and *π*/4) the tissue passively stretches resulting in σ_DV_ greater than the local value of σ_a_. If pre-stress is increased by a multiplicative factor, mimicking the increase of MyoII activity observed *in vivo,* these features are dramatically accentuated (Fig. 2d–f, Supplementary Fig. 1a and d–f).

The trace of the stress tensor, presented in Fig. 2b and d, is positive everywhere except at the anterior and posterior ends of the mesoderm region which are under net compressive stress in Fig. 2b. However, the principal stresses, which are the eigenvalues of the stress tensor, show a more complex pattern (Supplementary Fig. 1b and d). While both principal stresses are positive in the mesoderm, one of them is negative, denoting directional compression, just beyond the periphery of the mesoderm. The compressive stress is oriented orthogonally to directions pointing to the ventral furrow: it is thus parallel to the AP axis at lateral positions and parallel to the DV axis at anterior and posterior positions. This feature was not reported in 2D planar models [49, 50]. This compressive pattern can be intuitively understood by the fact that cells beyond the periphery of the mesoderm move centripetally towards the ventral midline, which generates a compressive hoop stress (Fig. 2e).

### 2.4 *In vivo* surface shape changes are reproduced by the mechanical model

In order to validate our mechanical model, we compared the dynamics of area change obtained computationally to experimentally measured changes in cell surface area while imposing a localized pre-stress increase proportional to MyoII intensity distribution measured experimentally in both space and time (Fig. 1g and 3a). MyoII activity increases strongly in the mesoderm as VFF engages. Quantitatively, the evolution of background-subtracted MyoII fluorescence in cells at the ventral midline is well approximated with a double-exponential function:

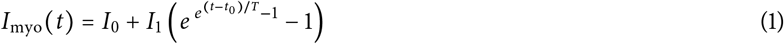

where *I*_0_ and *I*_1_ are coefficients in arbitrary units of fluorescence intensity, *t*_0_ the time from which MyoII shows a progressive increase, and *T* the characteristic time of increase of MyoII. The rate of these dynamics, reflected in the constant *T*, depends on temperature, however the time profile is highly reproducible for different experiments (Fig. 3a, see Materials and Methods). We fitted the pre-stress with MyoII temporal dynamics for an optimal match with the first part of the process of furrow formation (i.e., until minute 1 in Fig. 3a). In our computational model, as pre-stress gradually increases, the ventral region reduces in area showing similar dynamics as measured *in vivo* (Fig. 3a). *In vivo,* cells located close to the ventral midline exhibit an area reduction which is proportional to MyoII intensity increase. By contrast, cells located further away from the midline (between the 6th and the 9th cell row) increase in area (Fig. 3b) although their MyoII intensity also increases at a lesser rate (Fig. 1g). This corroborates previous experimental evidence and quantitative analysis showing that cells further away from the midline increase their surface area during furrow formation [9, 49, 4, 29, 52]. This spatial strain pattern is reproduced by our computational model (Fig. 3b). These results give rise to a seeming paradox: how can cells increase their surface area while simultaneously increasing the level of MyoII (i.e., the constricting pre-stress σ_a_)? To find an answer to this question we further investigated cell shape changes by quantifying the AP and DV lengths of cells located at different DV positions from the ventral midline. Cells with increasing surface area show a significant increase in DV length (Fig. 3c and d, Supplementary Fig. 2a) in agreement with anisotropic tissue strain shown in Fig. 2a. Thus, we considered whether this area increase was directly linked to the DV extension found in simulations in ventro-lateral locations. Figure 3c–f shows indeed that all cells more than 3 cell radii from the midline are generally stretched along the DV direction from an early time in gastrulation, in fair quantitative agreement with simulations (see also Supplementary Fig. 2a and Supplementary movie 3). Consistently with our model and in agreement with [52], cells for which neighbouring cells exert an extrinsic tensile stress larger than their own MyoII-based pre-stress are thus being stretched towards these neighbouring cells. The contraction along the AP direction, which is observed experimentally and which simulations predict (Supplementary Fig. 2a and c-e), is of smaller magnitude for cells farther from the midline, and therefore the combination of DV stretch and AP contraction results in an area increase for cells located on the ventro-lateral position.

**Figure 3:**
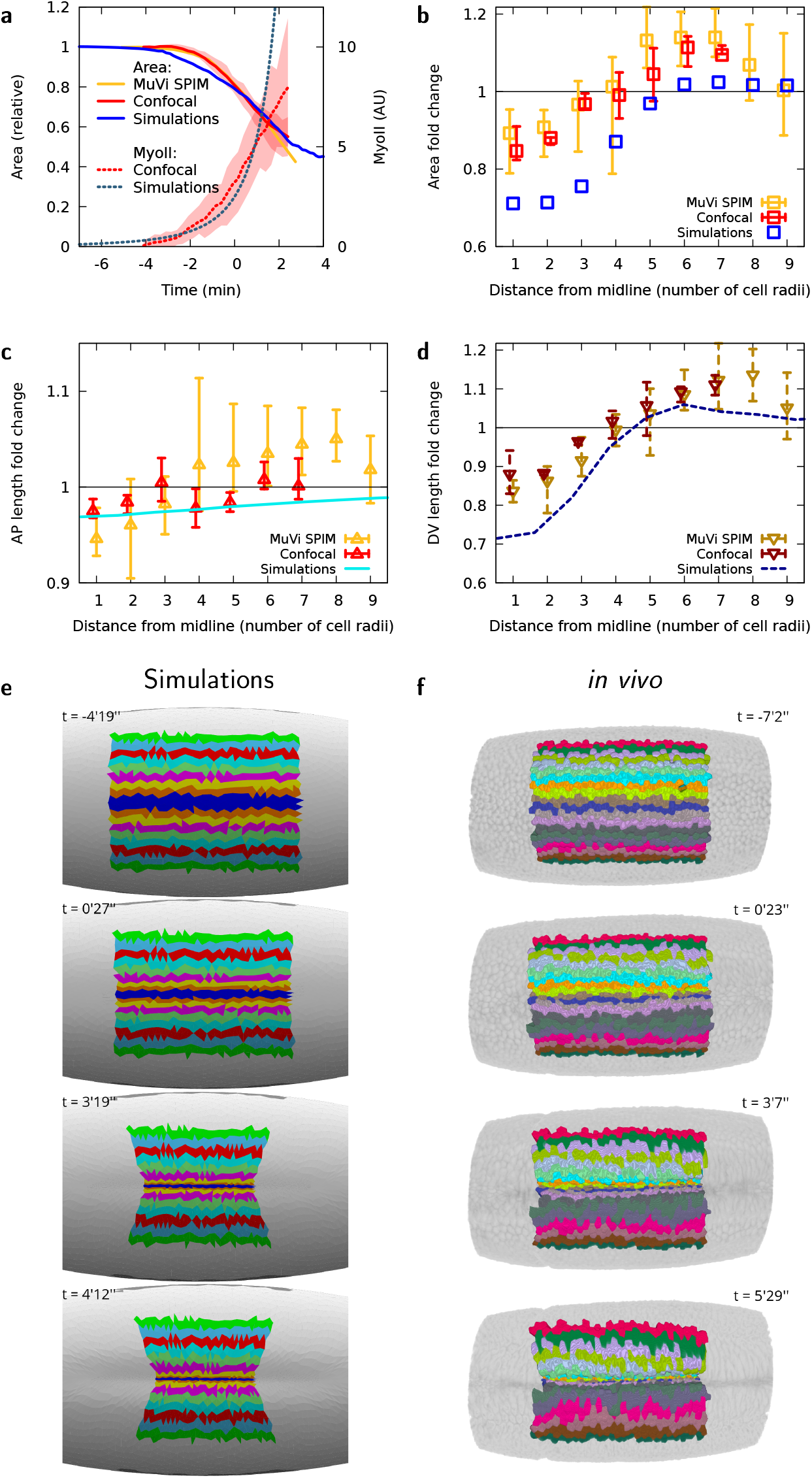
*In vivo* apical area changes are reproduced by the computational model. (**a**) MyoII average intensity and mesoderm apical area changes as a function of time, for *in vivo* analysis and simulations. (**b**) Apical area fold-change relative to the initial area of cells at different lateral distances from the midline at *t* = −1 min, in observations for *in vivo* analysis and simulations. (**c**) Apical AP size fold-change relative to the initial size for cells at different lateral distances from the midline at *t* = −1 min, (**d**) Apical DV size fold-change relative to the initial size for cells at different lateral distances from the midline at *t* = −1 min, for *in vivo* analysis and simulations. Panels **a-d,** 3 embryos using multi-view light sheet and 3 embryos using confocal microscopy, shaded areas in **a,** minimum and maximum values of confocal microscopy measurements, error bars in **b-d,** minimum and maximum values. (**e, f**) Time evolution of AP stripes of apical surface in simulation and MuVi SPIM, respectively.

### 2.5 Apical actomyosin mechanics drive AP midline flattening of the ventral tissue and furrow formation

Our model is now (i) tuned by imposing a pre-stress that is proportional to the experimentally measured MyoII distribution and (ii) validated since the imposed pre-stress results in surface area changes of the elastic sheet in good agreement with cell apical surface changes measured *in vivo.* Remarkably, the 2D elastic sheet forms a buckle resulting in a furrow along the long axis of the 3D ellipsoid in the region under pre-stress (Fig. 4a and Supplementary Fig. 3a). This shows that forces applied at the surface of an ellipsoidal 3D shape can be sufficient to drive the formation of a furrow. We then tested the predictive power of our model. We analyzed the furrow at different positions from the poles: the furrow forms first and is deeper in the mid region of the ellipsoid and appears with some delay in regions closer to the poles (Fig. 4b and c). We then measured furrow depth at different anterior-posterior positions *in vivo.* To that end, we embedded the embryo in a soft gel cylinder (preserving the embryo shape) and imaged it with multi-view light sheet microscopy (MuVi SPIM) to obtain isotropic resolved images (Materials and Methods). The embryo shows the same features as the computational model: the furrow forms deeper in the mid region of the embryo and eventually propagates towards the anterior and posterior poles (Fig. 4b, c and d, Supplementary movie 4), even though MyoII apical accumulation is uniform and does not form a propagating wave (Supplementary Fig. 3b). The curves representing the absolute furrow apex position at different AP locations in the embryo over time, eventually merge, highlighting a remarkable feature of VFF: while folds at distinct AP positions initially form at a different rate, eventually they align, sequentially reaching the same absolute apex position (Fig. 4b, Supplementary Fig. 3c and Supplementary movie 5). To better decipher the dynamics of ventral furrow propagation in the embryo, we digitally sectioned the 3D image of the embryo along the mid-sagittal plane (the plane separating the left from the right side of the embryo and intersecting the furrow midline, Supplementary Information). The mid-sagittal view of the embryo reveals a new feature: during furrow formation, the ventral tissue midline flattens along the embryo AP axis (Fig. 4e and Supplementary movie 4). Interestingly, the acellular embryo shows similar furrow formation features (Supplementary movie 6). While ventral flattening along DV has been characterized previously [8], AP flattening along the ventral tissue midline can only be seen in a mid-sagittal view which has not been reported before. To better characterize the dynamics of furrow formation in 3D, we measured and analyzed the changes of ventral tissue curvature along both the DV and AP tissue midlines (Fig. 4f, Supplementary Fig. 3d). Ventral curvature analysis for both our computational model and the embryo show the same trend: the curvature of the AP ventral midline gradually decreases eventually reaching the zero value (i.e., a flat tissue), concomitantly the ventral DV curvature decreases much faster until suddenly transitioning from positive to negative values (i.e., from convex to concave). Although this process is abrupt, it is not discontinuous (see Supplementary Fig. 3e) and it starts from a finite threshold value of pre-stress. Finally, this results in the formation of a furrow.

**Figure 4:**
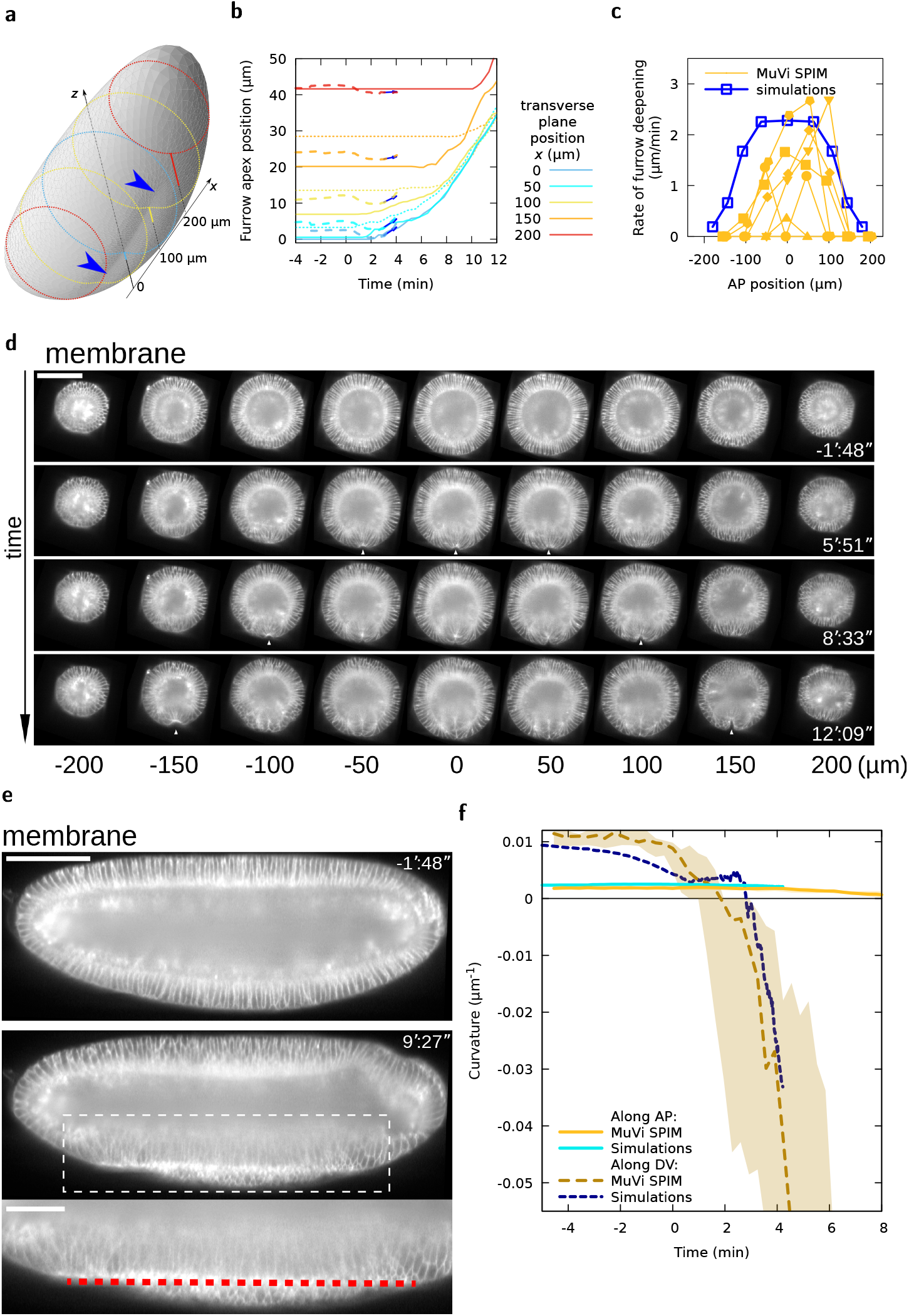
VFF results from tissue curvature changes along the DV and AP axes. (**a**) Embryo shape during VFF in simulations at *t* = 4’12”. Shading reveals furrow shape, blue arrowheads. Dotted lines are transverse cuts, solid lines give the furrow apex offset from a reference *z* position at different *x* positions. (**b**) Furrow apex position at different AP positions as a function of time in MuVi SPIM experiments (solid lines, posterior side, dotted lines, anterior side, *n* = 6) and simulations (dashed lines and arrow showing slope at *t* = 3’). (**c**) Rate of furrow formation at different AP positions at *t* = 3’. (**d**) Digital cross-sections at different AP positions. White arrowheads indicate VFF initiation. Scale bar 100 μm. (**e**) Digital mid-sagittal section of the embryo. Red line indicates ventral tissue flattening. Scale bars 100 μm, zoom 50 μm. (**f**) Curvature of the ventral tissue along the AP and DV axes as a function of time (MuVi SPIM, *n* = 6 embryos, shaded area denotes minimum and maximum).

### 2.6 The embryo anterior and posterior polar caps function as anchoring sites for the contracting ventral tissue

The ventral region of the ellipsoidal elastic sheet contracts under pre-stress (Fig. 2) resulting in sheet deformation. While along the DV axis the ventral sheet constricts in the mid region and stretches in more lateral regions, along the AP axis the sheet shows less deformation (Fig. 2a, Supplementary Fig. 1e, Fig. 3c–f). Since AP pre-stress drives little AP strain, it results in stress within the contractile region which can only be balanced by forces acting upon neighbouring tissues. We thus focused on the neighbouring tissue forming the polar caps. The polar tissues are submitted to both pressure forces exerted by the incompressible cytoplasm (Fig. 5b, gray arrowheads) and by pulling forces exerted by the contracting ventral sheet and which are close to the midline (Fig. 5b, red arrows, and d). During sheet contraction, the polar caps are pulled inwards leading to an increase of perivitelline space at the poles (Fig. 5a and b, dashed line). Remarkably, the same process occurs also *in vivo* (Fig. 5c) and tension is also maximal close to the midline (Fig. 5e). Therefore, the anterior and posterior polar caps may function as symmetrically positioned anchoring sites between which the ventral tissue midline flattens, working as a contractile string driving furrow formation. These ‘anchors’ are not immobile and result from the combination of the embryo geometry, internal pressure, and epithelial cohesion that provide resistance to the polar cap displacement towards the ventral region. To test if the position of anchoring sites could bias AP tissue midline flattening, we implemented IR fs laser cauterisation to establish ectopic anchoring sites (see [2, 53, 54, 4], and Materials and Methods). After generating two fixed sites at asymmetric positions along the AP axis of the embryo, the ventral tissue contracting in between the two ectopic fixed points and the AP midline still flattens, preserving tissue furrowing (Fig. 5f and Supplementary movie 7). Remarkably, under asymmetric boundary conditions, the ventral tissue midline now flattens along a direction that is no longer parallel to the AP axis but instead follows a line intersecting the two ectopic anchoring sites (Fig. 5f, dashed line). This shows that the position of anchoring sites defines the boundary conditions controlling the direction of AP tissue midline flattening during furrow formation.

**Figure 5:**
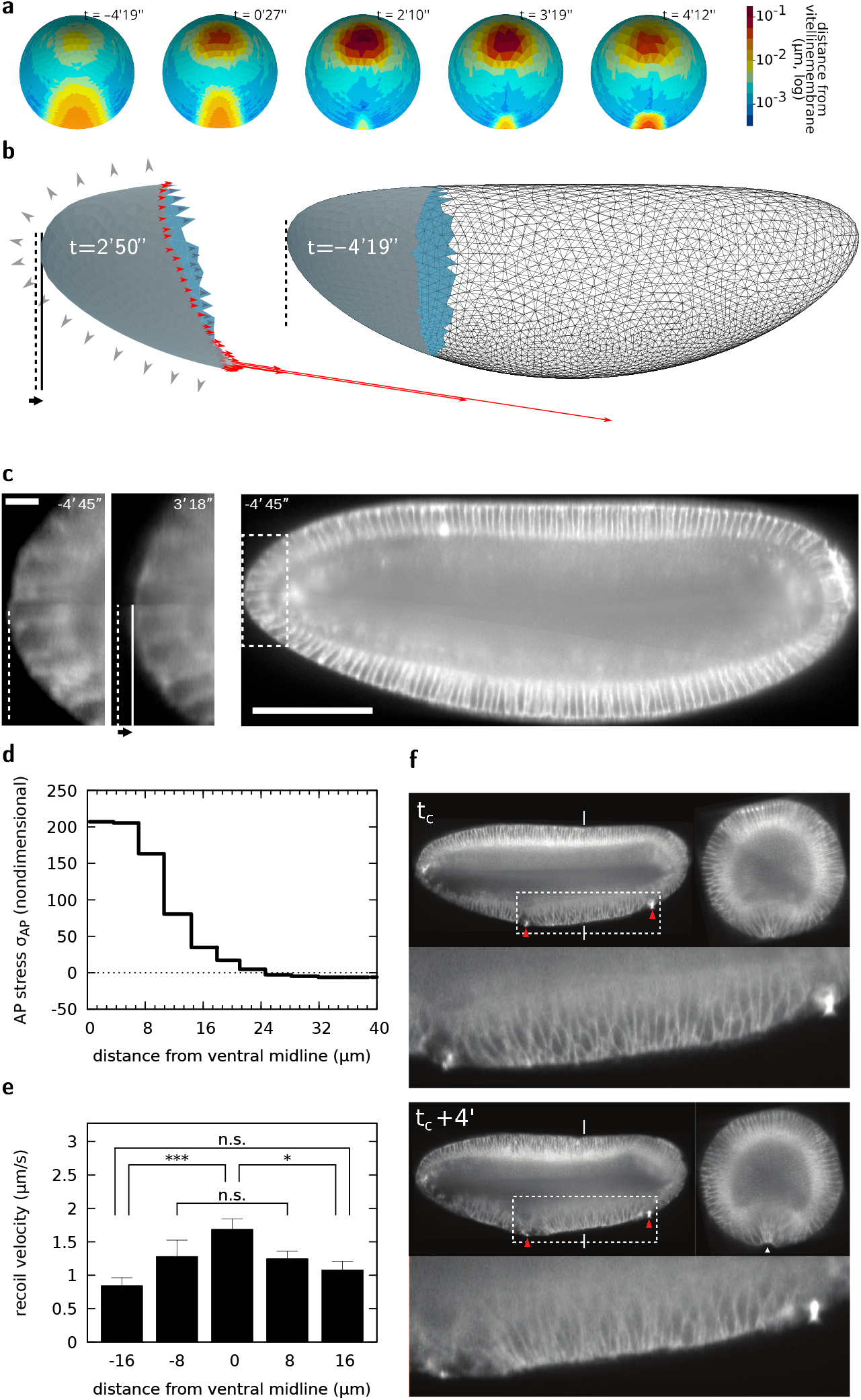
Embryo poles function as anchoring sites for ventral midline flattening and furrow formation. (**a**) Distance map of the apical surface to the vitelline membrane at different phases of VFF. (**b**) Forces exerted on the poles by the rest of the tissue (red arrows), and pressure forces exerted by the incompressible cytoplasms (gray arrowheads), pole tissue deformation (compare shape of solid-filled regions) and displacement of the pole (solid and dashed lines) in simulations. (**c**) Digital mid-sagittal section showing inward displacement of the pole tissue during VFF. Scale bar 100 μm, zoom 5 μm. (**d**) Tension distribution at different DV positions from the ventral midline in simulations at t=0’30”. (**e**) Recoil velocity distribution after DV-oriented IR fs laser ablation at different DV positions from the ventral midline. (**f**) Digital mid-sagittal and cross sections of an embryo on which two cauterizations (red arrowheads), acting as fixed points, have been performed at the ventral side (tc indicates time of cauterization). The red line indicates tissue straightening along the embryo mid-sagittal section in between the two cauterized regions. Experiment performed and result reproduced 3 times. Scale bar sagittal view 100 μm. Scale bar cross-section 50 μm.

## 3 Discussion

Tissue furrowing is a fundamental process during embryo gastrulation and neurulation. The mechanical process responsible for furrow formation is unclear. In this study we use as model system the *Drosophila* embryo and study furrow formation of the prospective mesoderm at the onset of gastrulation. While previous work has supported the idea that forces along the lateral side of cells play a key role and that cell wedging is a necessary step in furrow formation [19,13], we now challenged this view by developing a computational model based on a thin ellipsoidal elastic sheet in the 3D space, backed by 3D imaging, laser-based manipulation and multidimensional image analysis. Our model is based on two key assumptions imposed for simplicity: i) the sheet has homogeneous linear mechanical properties, with a Poisson modulus and a ratio of Young to bending moduli which are tuned to match observed in-plane deformations, and ii) the sheet is purely elastic, therefore, viscous properties are neglected. This last assumption, while not suited for modelling mesoderm internalization (during which mesoderm cells intercalate [55]), may be suitable to model the initial rapid process of mesoderm furrowing. By imposing a ventral pre-stress proportional to MyoII distribution measured *in vivo*, we show that our computational model can predict the magnitude and the dynamics of furrow formation and of cell apical shape changes. The concordance between our prediction and the observed dynamics confirms that the viscous response is negligible compared to the tissue elastic response, presumably because the mechanical load generated by the pre-stress is rapidly increasing and does not allow relaxation to occur. Importantly, our model shows that stress at the apical surface of a thin curved elastic sheet in the 3D space is sufficient to drive the formation of a furrow. After forming a furrow, the elastic sheet does not invaginate. This shows that apical mechanics, while being sufficient to drive the formation of a furrow, is insufficient for subsequent tissue internalization. These computational results are in agreement with the phenotype shown by acellular embryos that are able to form a ventral furrow but fail to internalize it. Therefore, cytoplasmic compartmentalization, cell lateral or basal forces and cell wedging, while dispensible for furrow formation, may be necessary for the second phase of mesoderm invagination during which the formed furrow is internalized.

Remarkably, the dynamics of furrow formation differ from those of MyoII and of cell shape change: while the change of MyoII intensity and cell apical area are smooth, the furrow forms abruptly (as shown by our DV curvature analysis). Such abrupt dynamics are reminiscent of the mechanical process of sheet buckling. At the onset of MyoII increase, the surface area of ventral cells is strongly reduced (from 20% at time 0 to 40% at 2 minutes). Our model shows that, during this first phase, most of the energy corresponding to the pre-stress results in DV contraction of ventral cells and in AP tension along the ventral midline. In this way, the DV curvature of the embryo surface at the ventral midline is reduced and eventually stalls at a smaller value (i.e., the tissue tends to flatten), whereas the AP curvature, which is initially smaller than the curvature along the DV axis, does not significantly change. This first phase is then followed by a second phase that results in an abrupt change of sign of the tissue curvature (i.e., from convex to concave) along the DV axis: this corresponds to the first appearance of the furrow that subsequently deepens over time. In our model, a new mechanical balance characterizes this second phase, during which most of the energy corresponding to the pre-stress is converted to work to bend the sheet. It is important to note that, even though the formation of the furrow is abrupt, there is no discontinuity in the sequence of tissue shapes. Since our model is free of energy dissipation, each of these shapes are at equilibrium. Thus, the control parameter (the tissue pre-stress reflecting MyoII activity) unequivocally determines the embryo surface profile. In plants, buckling has been shown to drive tissue deformation at a time scale much shorter than tissue growth [56,57]. In these cases the buckling dynamics are set by either inertial or viscous forces opposing the motion. In our model, the buckling dynamics are under the control of the active molecular mechanisms generating pre-stress during VFF.

Our model makes the prediction that the ventral furrow forms by propagating from the medial towards the polar regions of the embryo driven by a homogeneous distribution of MyoII along the ventral tissue. Our multidimensional *in vivo* analysis confirms furrow propagation in the absence of a MyoII wave. Furrow propagation emerges from the ventral tissue that folds sooner and faster in the central region of the mesoderm compared to regions closer to the poles, so that the furrow reaches the same absolute depth at different AP positions. This results in the flattening of the mesoderm at the ventral AP midline (i.e., at the furrow apex). Our study shows that the ventral midline, subject to the highest stress, works as a contractile string (like a ‘cheese cutter wire’) forming the furrow. The polar caps are pulled by the contracting ventral sheet, resisting the AP stress and therefore working as anchoring sites. The position of the polar caps imposes a ventral flattening that is parallel to the AP axis. Therefore, our study shows that embryo-scale mechanics, rather than local cell shape changes, is the key mechanism driving a buckling of the epithelial surface, forming a furrow which propagates and initiates mesoderm invagination. Models based on mesoderm cell-wedging would instead fail to reproduce furrow propagation in the absence of a MyoII wave.

Mechanical balance results from the interaction among the material units constituting the embryo. Since mechanical interactions span the entire embryo and are much faster (almost instantaneous) compared to the establishment of morphogen gradients or mechano-chemical waves, they account for long-range tissue interaction and eventual coordination. A 2D model of a 3D system, while potentially effective to provide some level of understanding with less computational resources and programming complexity, may bias the understanding or miss key aspects of a process that emerges from the global mechanical balance and the intrinsic 3D physical nature of the system. While in the past, the choice of performing 2D imaging, analysis and computational modelling was often dictated by technology limitations, we are now at an exciting time when we can study morphogenesis by performing multidimensional image analysis and by bridging scales from the embryo to the cell and vice versa. New imaging technology provides a synthetic view of the coordination of tissues at the scale of the whole embryo with subcellular resolution [1, 4, 3, 2] while computational modelling allows the real embryo geometry to be accounted for [5, 58, 7], opening new avenues to unravel the physical principles governing morphogenesis.

## 4 Methods

### Mutants and fly stocks

All fly stocks were reared on standard corn meal in a 25°C incubator. For collecting embryos for live imaging, flies were kept in cages and were allowed to deposit eggs on agar plates. *ubi*:Gap43::mCherry;; *klar* flies [55] that express a plasma-membrane targeted protein fused to mCherry fluorophore were used for 3D segmentation of mesoderm cells during folding. In all other experiments to monitor membranes in live embryos were collected from *ubi*:Gap43::mCherry (I) fly stock. For laser ablation experiments, embryos were collected from Sqh::mCherry; Spider::GFP fly stock, in which the Myosin II regulatory light chain (MRLC) (Spaghetti Squash, Sqh, in *Drosophila*) is labelled by mCherry and membrane by Spider (Gilgamesh) fused to GFP. For wild-type embryo cross-sections depicting MyoII localization, embryos were collected from fly stock expressing Sqh::mCherry. For cross-sections of acellular mutant embryos showing MyoII localization, embryos were collected from Δ*halo slam dunk*^1^/CyO, SqhGFP fly stock. *sqh^AX3^; sqh-GFP; GAP43-mCherry* fly strains were used for confocal imaging as described in [59].

### Actomyosin meshwork ablation

Apical actomyosin meshwork was ablated using a tunable femtosecond-pulsed infrared laser (IR fs, MaiTai) mounted on a Zeiss LSM 780 NLO confocal microscope tuned at 950 nm. Experiments were performed using a 40× 1.2 NA objective, and the bleach mode on the Zeiss Zen software (140 mW laser power at the focal plane, single iteration, 1 μs pixel dwell, 1 s frame rate). For segmented ablations and experiments to probe the necessity of apical actomyosin contractility for furrowing, only single ablations were performed. For experiments to block furrow invagination, a grid pattern of iterative ablations of the apical actomyosin network were performed every time the network was recovering.

### In toto embryo imaging and laser cauterization

Embryos were staged and dechorionated in bleach before mounting. Embryos were mounted in a glass capillary filled with 0.5% gelrite, with their long axis parallel to the capillary. A small portion of gelrite containing the embryo was then pushed out from the capillary. The embryo was imaged on a MuVi SPIM (Luxendo, Bruker) equipped with Olympus 20× 1.0 NA objectives and 488 nm and 594 nm lasers. Z-stacks were acquired with a step-size of 1 μm and during each acquisition, embryos were imaged in two opposing orthogonal views (0°-dorsal-ventral view, 90°-lateral view). Thus, for every single time point, four 3D stacks were recorded. Fusion of four stacks was obtained by using Matlab [4] resulting in a final isotropic pixel resolution of 0.29 μm. Laser cauterization was performed by coupling a femtosecond 920 nm laser (Alcor2, SPARK LASERS) to MuVi SPIM and by following a similar protocol as presented in [53].

### Confocal imaging and analysis

Apical shapes and *sqh*-GFP intensities were calculated for mesoderm cells during gastrulation for three wild-type embryos imaged and analysed in [59]. Embryo movies SG_1_, SG_3_ and SG_4_, which started prior to apical constriction and with similar overall *sqh*-GFP intensities, were used for this study. Confocal live imaging and cell tracking are as described in [59], with the evaluation of cell apical dimensions taking into account the local inclination of the embryo [60]. Mesoderm cells were identified as all cells that did not remain on the surface of the embryo after gastrulation. For every tracked mesoderm cell in each movie frame, 30 seconds apart, apical cell area, AP and DV lengths of the best-fit ellipse to the cell shape, distance of the cell centroid from the ventral mid-line and total apical *sqh*-GFP intensity were extracted.

### Time synchronisation

Time 0 was set as the instant when cells within the 5 rows closest to the midline had their area reduced by 20% compared to pre-gastrulation, allowing to synchronize all embryos, whether imaged with SPIM or confocal. Interpolation between datasets and time derivatives were calculated by LOESS moving window technique.The dynamics of development was found to be different between MuVi SPIM and confocal experiments due to differences in temperature. It was found that area reduction in confocal experiments was taking place at a twice slower rate than in MuVi SPIM, this factor was applied in Fig. 3a and for all subsequent confocal experiments. This corresponds to *T* = 20 min in Eq. 1.

### Numerical simulations

The geometry of the elastic surface used to construct the initial finite element mesh, as well as the volume constraint and constraint imposed by vitelline are decribed in Supplementary Information. A linear elastic model is then solved with the Surface Evolver software [46] with increasing pre-strain for the ventral region highlighted in Fig. 1f, as described in Supplementary Information.

## Acknowledgements

AJ, JE and MR acknowledge that this research was supported in part by the National Science Foundation under Grant No. NSF PHY-1748958 while in KITP at UC Santa Barbara. AJ, BD and MR thank the company Luxendo Bruker and SPARK LASERS for fruitful collaborations. We thank PRISM imaging facility for technical support. This work was supported by the French government through the UCAJEDI Investments for the Future project managed by the National Research Agency (ANR-15-IDEX-01), the Investments for the Future LABEX SIGNALIFE (ANR-11-LABX-0028-01), the ATIP-Avenir program of the CNRS, the Human Frontier Science Program (CDA00027/2017-C) and the Region SUD PACA Research and ERC Booster programs. GB, CL and BS acknowledge financial support of a Wellcome Trust Investigator Awards 099234/Z/12/Z and 207553/Z/17/Z. JF, AT, JE, CQ and PM acknowledge financial support from the European Community’s Seventh Framework Programme (FP7/2007-2013) ERC Grant Agreement Bubbleboost no. 614655, and are members of GDR 3570 MecaBio and GDR 2108 AQV of CNRS. JE was additionally supported by IRS “AnisoTiss” of Idex Univ. Grenoble Alpes. GB, CL, BS and JE benefited from a PICS CNRS travel grant and ANR-11-LABX-0030 ‘Tec21’ grant. The computations were performed using the Cactus cluster of the CIMENT infrastructure, supported by the Rhône-Alpes region (GRANT CPER07_13 CIRA). The authors thank Philippe Beys who manages the cluster, and K. Brakke for developing and maintaining the Surface Evolver software as well as invaluable interactions during this work.

## Author contributions

JE, PM, CQ and MR conceived the project. MR planned the *in vivo* experiments. BS planned the confocal experiments. JE, PM and CQ planned the modelling and computational approach. JF and CQ performed the computations. JF, CQ, PM and JE analysed the computational results. AJ, BD and MR performed the laser-based manipulations and light sheet experiments and analysis. GM provided support on the 3D image processing. CL and GB performed the analysis on confocal images. JE and MR wrote the manuscript. All authors commented on the manuscript.

## Supplementary Movies

Supplementary movie 1: Time-lapse showing actomyosin network recoil after IR fs laser dissection and eventual actomysin network recovery. Scale bar 10 μm.

Supplementary movie 2: Time-lapse showing mesoderm internalization failure as a consequence of periodic ventral tension inhibition. The ventral actomyosin network is periodically dissected using a IR fs laser over a grid patterned ROI. Scale bar 10 μm.

Supplementary movie 3: 3D rendering of segmented apical surfaces of ventral cells. Scale bar 50 μm.

Supplementary movie 4: Time-lapse showing digital sections along the sagittal plane (top) and along different cross-section planes at different AP positions in a wild-type embryo. Scale bar 100 μm.

Supplementary movie 5: Time-lapse showing different cross-section planes at different AP positions in a wild-type embryo and eye guide highlighting furrow apex position. Scale bar 100 μm.

Supplementary movie 6: Time-lapse showing digital sections along the sagittal plane (top) and along different cross-section planes at different AP positions in a slam-dunk-embryo. Scale bar 100 μm.

Supplementary movie 7: Time-lapse showing ventral tissue flattening along a line connecting cauterized loci. Scale bar 100 μm.

---

**Supplementary Figure 1:**
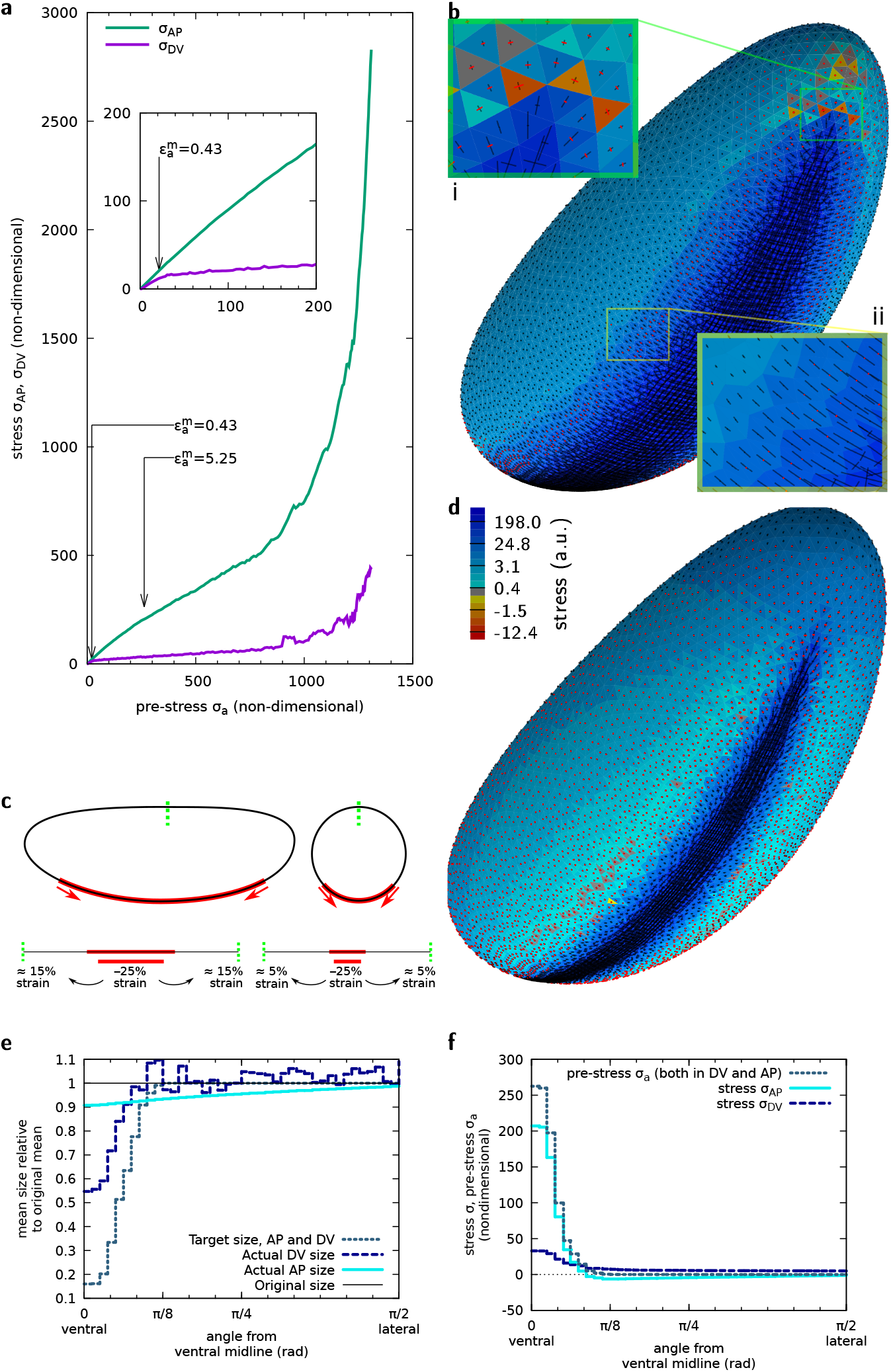
(**a**) Evolution of AP and DV stresses at the ventral midline as a function of the pre-stress σ_a_. (**b**) Principal stresses in each facet for midline pre-strain 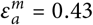. Black segments, positive principal stress (tensile); red segments, negative principal stress (compressive) along the corresponding directions. Facet colours, sum of the principal stresses, see **d** for colour code. Inserts (i) and (ii), 3× zoom of the regions outlined in white and in yellow. In (ii), tensor components are further enlarged by a factor 2. (**c**) Representation in sagittal and transverse sections of an example 25% strain in the ventral region and of the strain it implies in other regions if the embryo shape is unchanged. (**d**) Principal stresses in each facet for midline pre-strain 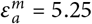. Same colour code as **b.** (**e**) Same as Fig. 2a but for 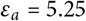. (**f**) Same as Fig. 2c but for *ε_a_* = 5.25.

**Supplementary Figure 2:**
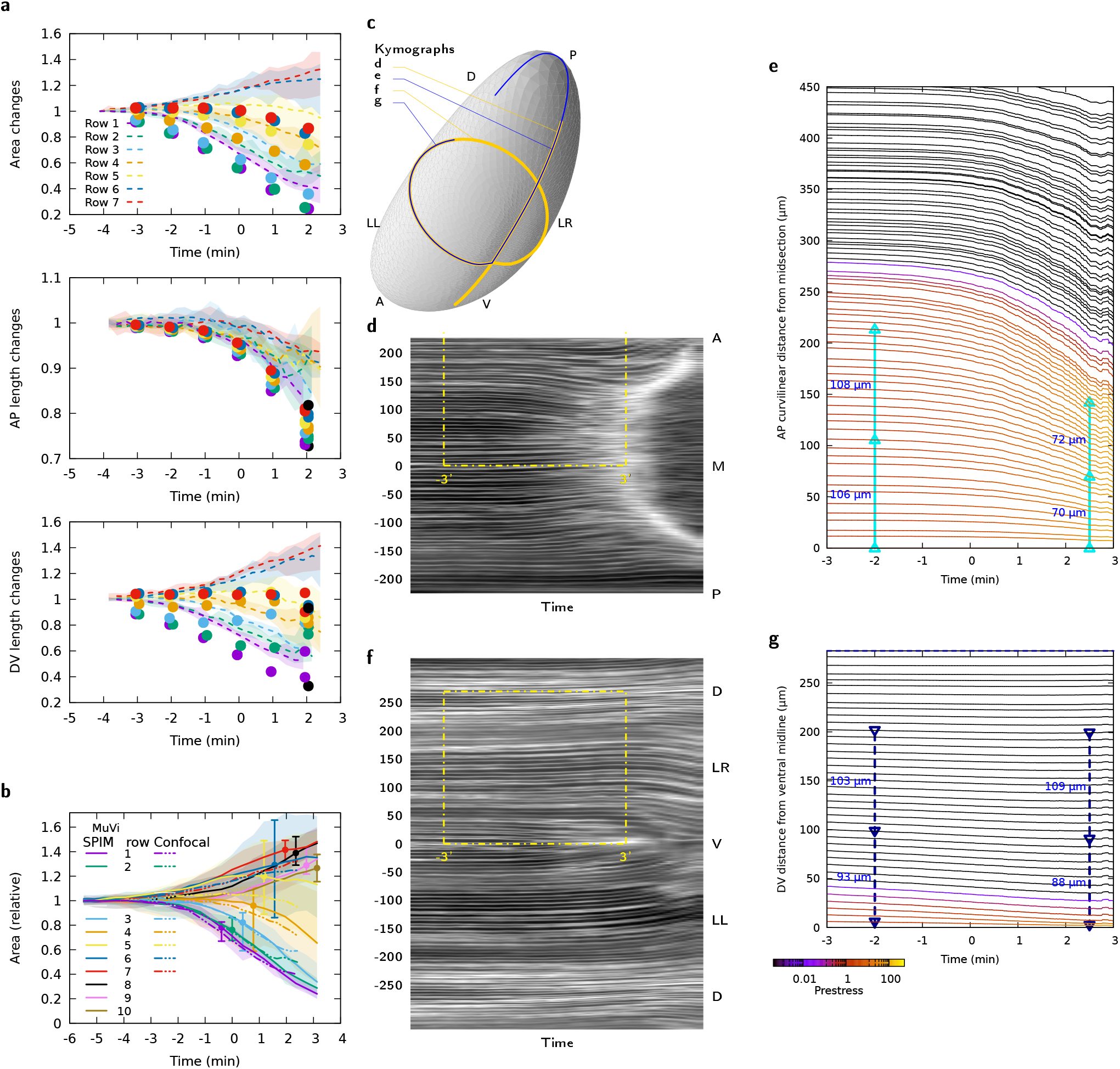
(**a**) Time evolution of apical area, AP and DV sizes of cells at different distances from the midline, in confocal experiments (lines) and simulations (symbols). (**b**) Time evolution of apical area of cells in MuVi SPIM and confocal experiments. (**c**) Localisation of kymographs of panels **d-g** on the apical surface of the embryo. (**d**) Kymograph of membrane signal in MuVi SPIM along AP (mid-sagittal line). M, mid; A, anterior; P, posterior, coordinates from mid-transverse plane in μm. Yellow box delineates the zone also shown in panel **e.** (**e**) Same as **d** in simulations, with colour coded pre-stress value (see colour bar in **g)** and quantification of the decrease of length between mid-transverse point and points marked with symbols. (**f**) Kymograph of membrane signal in MuVi SPIM along DV (mid-transverse line). V, ventral; LR, lateral right; LL, lateral left; D, dorsal, coordinates from the ventral midline in μm. (**g**) Same as f in simulations.

**Supplementary Figure 3:**
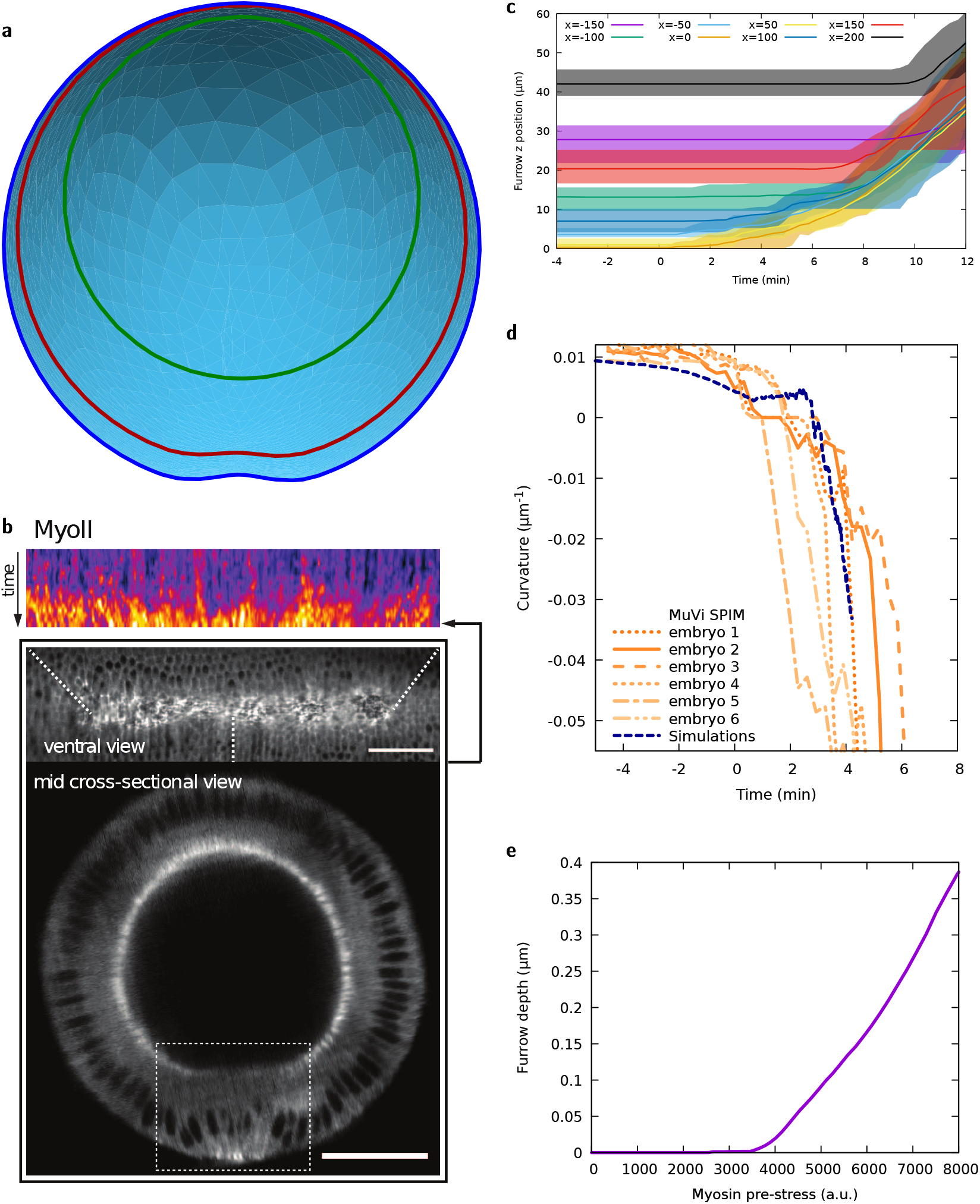
(**a**) See-through view of embryo shape in simulations at *t* = 4’12” from the posterior end. (**b**) Top, kymograph showing MyoII accumulation along the AP axis. Mid and bottom, ventral and mid-transverse view of the embryo before folding, respectively. Scale bars 50 μm. (**c**) Ventral furrow apex position at different AP positions. Lines represent medians and shaded areas the interval between min and max values. The analysis was performed on 6 embryos. Negative and positive *x* values indicate more anterior and more posterior positions, respectively. (**d**) Curvature of the ventral tissue along the DV axis as a function of time, with individual data from *n* = 6 MuVi SPIM embryos. (**e**) Depth of the ventral furrow as a function of the pre-stress *σ_a_* applied.

