## Supplementary Text for "Embryo-scale epithelial buckling forms a propagating furrow that initiates gastrulation"

### Quantifications in MuVi SPIM data

#### Measuring furrow propagation along the AP axis

Cross-sections were digitally made 50 μm apart along the AP axis. Furrow depth at different AP positions was measured using a dedicated point-picker ImageJ macro.

#### Measuring anterior pole distance

Mid-sagittal sections were digitally extracted from the time-lapses and the position of the vitelline membrane at the anterior pole and the apical position of the anterior-most blastoderm cell were recorded using the point-picker plugin. The relative apical position of the anterior-most blastoderm cell was calculated with respect to the position of the vitelline membrane at the anterior pole and this was plotted over time.

#### Measuring tissue shortening

Digital mid-cross and sagittal sections were obtained using ImageJ to measure tissue shortening along the DV and AP axes. Different embryos were aligned in time by keeping t0 as the frame at which cell apical area reduces to 20%. To measure tissue shortening along the AP axis, an identity cell was fixed at the half-length along the mid-sagittal section. Cells located 100 μm anteriorly and posteriorly from this point were marked and their distance from the identity cell was measured over time along the embryo surface. To measure tissue shortening along the DV axis, an identity cell was fixed at the midline of cross-sectional images at the ventral side. The distance along the embryo surface of the two cells located on opposite sides four cells away from the identity cell was then measured over time.

#### Recoil velocity

To measure actomyosin network recoil after ablation, the point-picker plugin was used to follow the cut end. Distance moved by the network after ablation was plotted against time and the maximum recoil velocity was measured by calculating the first derivative.

#### Curvature analysis

An ImageJ macro was developed to measure the radius (R) of a circle generated from three consecutive points. The curvature was calculated as 1/R and was plotted against time. Curvature of the mesoderm tissue was measured both along the AP and DV axes. A convex curvature is given by a positive and a concave curvature is given by a negative value.

#### 3D image segmentation and analysis

Mesodermal cells were segmented and tracked by inter-registration based on iterative projections of segmentations from one time point to another using the ASTEC algorithm (1). Morphological data were extracted and analyzed using Python. ImageJ dedicated macros were used for image treatment and the 3D Viewer plugin for rendering (2).

### Numerical simulations

#### Geometry of the elastic surface of the model

Following (3), we approximate the initial embryo shape by a closed surface of equation:

$$\Gamma_0 = \left\{ \left(\frac{x}{R_{AP}}\right)^2 + \left(\frac{y}{R_{DV}}\right)^2 + \left(\frac{z}{R_{DV}} - Z_p\left(\frac{x}{R_{AP}}\right)^2\right)^2 = 1 \right\}$$

This describes a prolate ellipsoid, with circular cross-sections. The mid-cross-section at $x = 0$ is a circle of radius $R_{DV}$ centered at $(x, y, z) = (0, 0, 0)$. For a positive parameter $Z_p$, this ellipsoid is bent with the center of cross-sections offset towards increasing $z$ as $|x|$ increases. In particular, the poles along the long axis are at the positions $(x, y, z) = (\pm R_{AP}, 0, Z_p)$.

As in (3), we choose $R_{AP} = 3R_{DV}$ and $Z_p = 0.45$, which defines the shape corresponding to the initial equilibrium configuration of the elastic surface in terms of in-plane stress. While all calculations are done in nondimensional units of $R_{DV}$, the dimensional counterparts of all quantities are then calculated with $R_{DV} = 90$ µm.

The elastic surface $\Gamma(\sigma_a)$ is then defined as the surface of minimal elastic energy for a given pre-stress field $\sigma_a$. The spatial dependence of $\sigma_a$ is described in the main text, the corresponding rheology is described in what follows.

Additionally, two constraints are applied to the configuration of the elastic surface $\Gamma$. First, it encloses a constant volume independent of $\sigma_a$ and equal to the initial volume of $\Gamma_0$. This models the fact that the permeability of the apical surface of the epithelium is low, such that water fluxes across it are assumed to be zero during the process of initial furrow formation. This gives rise to pressure forces which are dependent on $\sigma_a$ and are of uniform magnitude in space. Second, $\Gamma$ is itself enclosed in an undeformable vitelline membrane. The vitelline membrane is defined as a closed surface $\Gamma_V$ parallel to the initial elastic surface $\Gamma_0$, at a distance $2.5 \times 10^{-3} R_{DV} \simeq 0.2$ µm towards the exterior. Non-interpenetration of the elastic surface with the vitelline membrane is simulated by Surface Evolver constraint algorithm (4), a penalty method based on a level set function ensuring that the elastic surface is in the interior side of $\Gamma_V$.

#### Elastic model

The deformation of an elastic layer of non-vanishing thickness can be modelled using a surface with two local additive contributions: bending and in-plane deformation (5, 6).

The bending surface energy density writes $\frac{1}{2}\kappa\,(c - c_0)^2$ where $\kappa$ is the bending modulus of the surface, $c = \frac{1}{R_1} + \frac{1}{R_2}$ its mean curvature ($R_i$ are the principal algebraic curvature radii), and $c_0$ the (local) intrinsic curvature, taken to be the initial one in this work (5, 7).

We now proceed to describe the part of the surface energy density due to in-plane deformation. In a continuum description of in-plane deformations, the gradient deformation matrix $\mathbf{F}^c$, with $F^c_{ij} = \frac{\partial X_i}{\partial x_j}$ where $(X_1, X_2)$, describes the deformed position of a material point initially at the $(x_1, x_2)$ position (the indices stand for any two coordinates describing a surface), allow to define the right Cauchy-Green deformation tensor $(\mathbf{F}^c)^T\mathbf{F}^c$ and the Lagrangian finite strain tensor $\boldsymbol{\epsilon} = \left((\mathbf{F}^c)^T\mathbf{F}^c - \mathbf{I}\right)/2$, a widely used measure of how a material piece of surface locally differs before and after deformation.

The elasticity of the surface is described by the Hookean model, where the (locally) in-plane stress tensor $\boldsymbol{\sigma}$ is linked to the (locally) in-plane strain tensor $\boldsymbol{\epsilon}$. The model surface is considered as isotropic in plane, which allows to describe full elasticity using only two parameters, the 2D Young modulus $Y_{2\mathrm{D}}$ and the 2D Poisson ratio $\nu_{2\mathrm{D}}$. The surface energy density of in-plane deformation in this model can be expressed in two different useful forms:

$$\begin{aligned} e_{def} &= \frac{Y_{2\mathrm{D}}}{2(1+\nu_{2\mathrm{D}})}\left[\mathrm{Tr}(\boldsymbol{\epsilon}^2) + \frac{\nu_{2\mathrm{D}}}{1-\nu_{2\mathrm{D}}}(\mathrm{Tr}\boldsymbol{\epsilon})^2\right] \\ &= \frac{\chi_{2\mathrm{D}}}{2}(\mathrm{Tr}\boldsymbol{\epsilon})^2 + \mu_{2\mathrm{D}}\left(\mathrm{Tr}(\boldsymbol{\epsilon}^2) - \frac{1}{2}(\mathrm{Tr}\boldsymbol{\epsilon})^2\right) \end{aligned} \tag{1}$$

where $\chi_{2\mathrm{D}}$ and $\mu_{2\mathrm{D}}$ are, respectively, the 2D compression and shear moduli:

$$\chi_{2\mathrm{D}} = \frac{Y_{2\mathrm{D}}}{2(1-\nu_{2\mathrm{D}})} \qquad \mu_{2\mathrm{D}} = \frac{Y_{2\mathrm{D}}}{2(1+\nu_{2\mathrm{D}})}$$

In-plane stresses write:

$$\boldsymbol{\sigma} = \chi_{2\mathrm{D}}(\mathrm{Tr}\boldsymbol{\epsilon})\mathbf{P} + 2\mu_{2\mathrm{D}}\left(\boldsymbol{\epsilon} - \frac{1}{2}(\mathrm{Tr}\boldsymbol{\epsilon})\mathbf{P}\right)$$

with $\mathbf{P} = \mathbf{I} - \boldsymbol{n}\otimes\boldsymbol{n}$ the projection tensor along the surface.

We nondimensionalise the parameters with $\tilde{\chi}_{2\mathrm{D}} = \chi_{2\mathrm{D}}R^2_{DV}/\kappa$ and $\tilde{\mu}_{2\mathrm{D}} = \mu_{2\mathrm{D}}R^2_{DV}/\kappa$. All figures are shown for the choices $\tilde{\chi}_{2\mathrm{D}} = 50$ and $\nu_{2\mathrm{D}} = 0$.

### Numerical approach

We describe here the approach generally used in the Surface Evolver (4). Although these implementation details are not specific to our approach, they are necessary to introduce the numerical technique for simulation pre-strain in the next paragraph.

In a finite element description, $\mathbf{F}^c$ is approximated locally by $\mathbf{F}$, the matrix of the linear transformation from the unstrained to the strained facet, written in a local basis. Let $\mathbf{S} = [S_1, S_2]$ be the $2\times 2$ matrix formed by any 2 sides of the unstrained facet ABC,

$$\mathbf{S} = \begin{bmatrix} x_B - x_A & x_C - x_A \\ y_B - y_A & y_C - y_A \end{bmatrix},$$

and $\mathbf{W}$ the corresponding matrix of the same facet in the deformed configuration. Then $\mathbf{W} = \mathbf{FS}$ which rewrites $\mathbf{F} = \mathbf{WS}^{-1}$. With $\mathbf{F}^T = \left(\mathbf{S}^{-1}\right)^T\mathbf{W}^T$, one gets the numerical implementation for approximating the Lagrangian finite strain tensor in each facet of the finite element mesh:

$$\mathbf{E} = \frac{1}{2}\left(\left(\mathbf{S}^{-1}\right)^T\mathbf{W}^T\mathbf{W}\mathbf{S}^{-1} - I\right)$$

Note that this expression is invariant with respect to permutations between $A$, $B$ and $C$.

In practice, since $\boldsymbol{\epsilon}$ appears in Eq. 1 only through $\mathrm{Tr}\left(\boldsymbol{\epsilon}^2\right)$ and $\mathrm{Tr}\,\boldsymbol{\epsilon}$, one can use an alternative representation of the strain, namely $\mathbf{E}^* = \left(\mathbf{S}^T\right)^{-1}\mathbf{E}\mathbf{S}^T$, because due to matrix properties, $\mathrm{Tr}\,\mathbf{E}^* = \mathrm{Tr}\,\mathbf{E}$ and $\mathrm{Tr}\left(\mathbf{E}^{*2}\right) = \mathrm{Tr}\left(\mathbf{E}^2\right)$.

This definition is equivalent to $\mathbf{E}^* = \frac{1}{2}\left(\mathbf{W}^T\mathbf{W}\mathbf{S}^{-1}\left(\mathbf{S}^{-1}\right)^T - \mathbf{I}\right)$, or $\mathbf{E}^* = \frac{1}{2}\left(\mathbf{G}_W\left(\mathbf{G}_S\right)^{-1} - \mathbf{I}\right)$, where $\mathbf{G}_W = \mathbf{W}^T\mathbf{W}$ and $\mathbf{G}_S = \mathbf{S}^T\mathbf{S}$ are the (real and symmetric, hence diagonalisable) Gram matrices of, respectively, the strained and unstrained facets. Since Gram matrices, that make only intervene the square lengths and the scalar product of the two sides choosen to determine the facets, are easy to calculate from side lengths only, the alternative definition $E^*$ is preferred in the minimisation algorithm used for the equilibrium situations (4).

#### Pre-strain and pre-stress

Pre-strain corresponds to a reduction of the equilibrium configuration area. In order to apply pre-strain in a given facet, we multiplied the Gram matrix of the unstrained facet by a factor $\gamma < 1$ (which amounts to multiply the dimensions of the equilibrium configuration of the facet by $\sqrt{\gamma}$):

$$\mathbf{G}_S^\gamma = \gamma\mathbf{G}_S$$

This approach can be compared to morphoelasticity, see (8).

The alternative deformation tensor hence becomes:

$$\begin{aligned}\mathbf{E}_\gamma^* &= \frac{1}{2}\left(\frac{1}{\gamma}\mathbf{G}_W\left(\mathbf{G}_S^\gamma\right)^{-1} - \mathbf{I}\right)\\ &= \frac{1}{\gamma}\mathbf{E} - \frac{\gamma-1}{2\gamma}I\end{aligned}$$

The trace is easily calculated:

$$\mathrm{Tr}(\mathbf{E}_\gamma^*) = \frac{1}{\gamma}\mathrm{Tr}\mathbf{E} - \frac{(\gamma-1)}{\gamma}$$

The stress thus becomes:

$$\begin{aligned}\sigma_\gamma &= \chi_{2D}(\mathrm{Tr}\mathbf{E}_\gamma^*)I + 2\mu_{2D}\left(\mathbf{E}_\gamma^* - \frac{1}{2}(\mathrm{Tr}\mathbf{E}_\gamma^*)\mathbf{I}\right)\\ &= \chi_{2D}\left[\frac{1}{\gamma}\mathrm{Tr}\mathbf{E} - \frac{(\gamma-1)}{\gamma}\right]\mathbf{I} + 2\mu_{2D}\left(\left[\frac{1}{\gamma}\mathbf{E} - \frac{\gamma-1}{2\gamma}\mathbf{I}\right] - \frac{1}{2}\left[\frac{1}{\gamma}\mathrm{Tr}\mathbf{E} - \frac{(\gamma-1)}{\gamma}\right]\mathbf{I}\right)\\ &= \frac{\chi_{2D}}{\gamma}\left[\mathrm{Tr}\mathbf{E} - (\gamma-1)\right]\mathbf{I} + 2\frac{\mu_{2D}}{\gamma}\left(\mathbf{E} - \frac{1}{2}(\mathrm{Tr}\mathbf{E})\mathbf{I}\right)\end{aligned}$$

Area pre-strain is thus equivalent to an isotropic pre-stress $\sigma_a = \chi_{2D}\frac{\gamma-1}{\gamma}$. Note that the apparent elastic moduli of the material are also increased by a factor $1/\gamma$, due to the fact that their equilibrium configuration is of dimensions $1/\gamma$-fold smaller than their intial configuration, with respect to which strain is calculated.
